# Retinoid X Receptor as a Therapeutic Target to Treat Neurological Disorders Associated with *α*-Synucleinopathy

**DOI:** 10.1101/2024.06.10.598149

**Authors:** Assylbek Zhylkibayev, Christopher R. Starr, M. Iqbal Hossain, Sandeep K Barodia, Shaida A. Andrabi, Maria B. Grant, Venkatram R. Atigadda, Marina S. Gorbatyuk, Oleg S. Gorbatyuk

**Author notes:** Correspondence (O.S.G.); (M.S.G.). (A.Z.). (C.R.S.); (M.B.G.). (M.I.H.); (S.A.A.).

## Abstract

This study investigated the therapeutic potential of the nuclear retinoid X receptor (RXR) in mitigating the progression of alpha-synucleinopathies (αSNPs), particularly in Parkinson’s disease (PD). PD-like pathology in mice was successfully induced through the co-delivery of AAV expressing human α-synuclein (αS) and αS preformed fibrils (PFFs) into the substantia nigra pars compacta (SNpc). Significant increases in Lewy body (LB)-like inclusions, loss of tyrosine hydroxylase-positive (TH+) neurons, and reductions in dopamine (DA) levels in the striatum were observed. Additionally, diminished levels of PPARα and NURR1, along with elevated GFAP and Iba1, markers of neuroinflammation, microglial activation, and astrocytic gliosis were associated with PD pathogenesis. AAV-mediated overexpression of human RXRα demonstrated preservation of TH+ neurons, prevention of DA decline and attenuation of αS accumulation. Furthermore, RXR-treated PD brains showed a reduced number of GFAP+ and Iba1+ cells, decreased GFAP+ and Iba1+ immunoreactivity, and fewer and less widespread LB-like aggregates. RXR overexpression also enhanced the production of PPARα and NURR1, proteins critical for neuronal survival. These findings suggest that RXRα activation promotes neuroprotection by mitigating αSNPs and chronic neuroinflammation, a major contributor to PD progression. This research underscores the therapeutic potential of targeting nuclear receptors, such as RXR, in neurodegenerative diseases like PD

## 1. Introduction

Alpha-synucleinopathies (αSNPs) including Parkinson’s disease (PD) and dementia with Lewy bodies (LB), rank as the most prevalent neurodegenerative condition after Alzheimer’s disease (AD) [1]. These conditions are characterized by abnormal intraneuronal inclusions of α-synuclein (αS) [2]. While the pathology of PD is multifaceted, involving several interconnected processes, including the degeneration of dopamine (DA) nigral neurons, LB formation, mitochondrial dysfunction, protein aggregation, impaired clearance, and oxidative stress, chronic neuroinflammation stands out as a hallmark of PD pathophysiology [2,3]. The glial cell activation and heightened levels of pro-inflammatory factors constitute prevalent features of the PD-afflicted brain [4]. To date, only a few treatments exist that interfere with inflammation and immune deficiency, including minocycline, dexamethasone, and several non-steroid anti-inflammatory drugs. While these approaches protect DA neurons from degeneration and glial cell activation, a breakthrough therapy based on innovative molecular mechanisms and targets is an unmet need.

The modulation of nuclear receptors (NRs) has been proposed as one of the most attractive therapeutic strategies for correcting PD pathobiology [5-11]. The nuclear retinoid X receptor (RXR) is of particular interest for therapeutic intervention, due to its binding and activation of both permissive partners, NURR1 and PPARs, whose dysfunctions have been observed in AD, multiple-system atrophy, and PD [12-14] and which are known to regulate expression of inflammation-associated genes [15,16]. Despite the importance of pharmacological RXR activation against PD-causing toxins [5,17,18], effective RXR-based treatments for PD patients are still in development.

Cumulative evidence indicates that abnormal RXR signaling triggers neuronal stress and neuroinflammation in PD pathobiology [6,8]. Activation of RXR binding partners, either NURR1 or PPAR is beneficial for reversing the PD pathology seen in animal and cellular models of PD [5,7,11,15,19-21]. Furthermore, it has been shown that activation of RXR by treatment with the RXR ligand LG100268 protects cultured DA neurons from stress and PD-like pathology induced by toxin 6-hydroxy dopamine (6-OHDA) [6]. The activation of PPAR by the application of fenofibrate and GW501516 reduces neuroinflammation and PD neurodegeneration [15]. Activation of the NURR1/RXRα heterodimer by NURR1 agonists can stand as a monotherapy for PD to increase striatal DA levels and up-regulate NURR1 target genes in PD-like brain conditions [22]. Collectively, these studies lead us to ask whether RXR/PPAR and RXR/NURR1 complexes are destabilized in the experimental PD model and whether delivery of exogenous RXR to the PD brain activates PPARs and/or NURR1, providing an effective interventional treatment aimed at controlling inflammatory response and reversing PD pathobiology.

The absence of evidence for the direct effects of RXR up-regulation on αS induced neurodegeneration in animal models of PD and the disease related αSNPs fueled our interest in conducting the current study. Using recombinant adeno-associate virus (AAV) to overexpress human RXRα (AAV-RXR) in the mouse substantia nigra pars compacta (SNpc), we examined the role of RXR in PD pathogenesis. Specifically, we investigated the impact of RXRα on the inflammatory response and the changes in viability of nigral tyrosine hydroxylase positive (TH+) cells in response to αS-induced neurotoxicity.

## 2. Materials and Methods

### 2.1 Primary mouse cortical neuron culture

All the experiments involving the use of animals were approved by the Institutional Animal Care and Use Committee (IACUC) at the University of Alabama at Birmingham. Primary mouse embryonic cortical cells were isolated at embryonic day 15 (E15) as described previously [23]. Briefly, a day before isolation 12-well plates and glass slides were treated with attachment factor – poly-l-Ornithine (#P4957, Sigma-Aldrich, St. Louis, MO, USA). Mouse brains were dissected and placed in DMEM (#11-965-084, Fisher Scientific, Waltham, MA, USA) supplemented with 20% fetal bovine serum (#A4736401, Fisher Scientific, Waltham, MA, USA) were used as dissection medium. Cortical tissue was trypsinized using TypLE (#12-605-010, Fisher Scientific, Waltham, MA, USA) for 10 min at 37°C. Digested tissue was homogenized by pipetting in DMEM+10% fetal bovine serum and centrifuged at 1200g for 5 min. Supernatant was discarded and replaced with Neurobasal medium (#A1371001, Fisher Scientific, Waltham, MA, USA) containing B27 supplements (#A3582801, Fisher Scientific, Waltham, MA, USA). Tissue pellets were filtered through 40μm cell strainer (#50-146-1426, Fisher Scientific, Waltham, MA, USA) and transferred to plates and slides. On the second in vitro day (DIV 2), growth of glial cells was inhibited by adding 50 μM 5-fluoro-2-deoxyuridine (#f0503, Sigma-Aldrich, St. Louis, MO, USA) for 3 days. AAV-GFP or AAV-RXR were added at DIV-2 as well, while recombinant alpha synuclein pre-formed fibrils (PFFs) from Stress Marq Biosciences (#SPR-322, Victoria, BC, Canada) were added starting DIV 5 at a concentration of 5μg/ml and alpha synuclein protein from Stress Marq Biosciences (#SPR-322E, Victoria, BC, Canada) at a concentration of 5μg/ml. Cells were grown until DIV 14 and further harvested for analysis.

### 2.2 AAV vectors

All vectors were packaged in AAV2/5 capsid and purified as described previously [24]. Titers for all AAV constructs were equalized to 6 x10^11^ vg/ml.

### 2.3 Intracerebral injection of PFFs and AAV vectors

All surgical procedures were performed using aseptic techniques and isoflurane gas anesthesia, as previously described [25]. Stereotaxic coordinates for intranigral injections in C57BL6 mice (three-month-old males, 28±3 g) were AP -2.9 mm, ML +1.2 mm, and DV -4.1 mm, relative to the bregma. The total injection volume was 2.0 μl at a speed of 0.5 μl/min. For these injections, we used a pulled glass pipet with an outer diameter not exceeding 40–45 μm to avoid needle-related damage and reduce the injury-mediated inflammatory response.

### 2.4 Behavioral test

The cylinder test was employed to measure spontaneous forelimb use [26]. A mouse was placed in a glass cylinder and the number of times it rears up and touches the cylinder wall was measured. The wall touches were subsequently scored for the left and right paws. The data is expressed as the percentage of contralateral (left) to injected side paw usage relative to the total number of touches by both paws.

### 2.5 Isolation and Processing of Tissues

Animals were deeply anesthetized by ketamine (50mg/kg) and xylazine (10mg/kg) cocktail intraperitoneally. The brains were removed and divided into two parts. The caudal part containing the SNpc was fixed in ice-cold 4% paraformaldehyde in 0.1 M phosphate buffer, pH 7.4. The fixed parts of brains were stored overnight at 4°C and then transferred into 30% sucrose in 0.1 M PB for cryoprotection. Thirty μm thick coronal sections were cut and further processed for immunohistochemistry. The rostral piece of brain tissue was immediately used to dissect the right and left striatum. Tissue samples were frozen separately on dry ice and kept at -80° C until assayed. In this study, the mice from the 4-week time point of post-surgery were used to obtain SNpc tissue samples for Western blot analysis. Frozen brains were sectioned on a Leica CM1510S cryostat (Leica, Buffalo Grove, IL, USA) into 150 μm slices with SNpc tissue subsequently dissected out under a microscope, as described previously [24].

### 2.6 Immunohistochemistry

For the bright-field microscopy analysis, 30μm floating sections were preincubated with 1% H2O2–10% methanol for 15 min and then with 5% normal goat serum for 1 h. Sections were incubated overnight at room temperature with anti-TH (1:2000; #MAB318, mouse; Millipore) antibody. Next, incubation with biotinylated secondary anti-mouse antibody was followed by incubation with avidin–biotin–peroxidase complex (VECTASTAIN® Elite® ABC-HRP Kit, Peroxidase (Standard) (PK-6100) Vector Laboratories, Burlingame, CA, USA). Reactions were visualized using NovaRED Peroxidase (HRP) Substrate Kit (Vector NovaRED® Substrate Kit, Peroxidase (SK-4800), Vector Laboratories, Burlingame, CA, USA).

For confocal microscopy, sections were incubated with primary antibodies against human αS (1:1000; 32-8100, mouse; Invitrogen), pS129 (alpha-synuclein (phospho S129); 1:500; [EP1536Y] (ab51253); rabbit monoclonal; Abcam), and TH (anti-tyrosine hydroxylase antibody; 1:1000; AB1542, sheep; Millipore), human RXRα (RXRA monoclonal antibody (K8508); 1:200; 43-390-0; mouse , Invitrogen), and secondary fluorescent antibodies labeled with Alexa Fluor 488, 555, and 647 (1:500 for all; PIA32773-mouse; A21202-mouse; PIA32731TR-rabbit; PIA32794-rabbit; A21448-sheep, Invitrogen). The sections were examined using an AX-R Confocal Microscope coupled to a fully automated Ti2-E system (Nikon Instruments inc., Melville, NY, USA). NIS-Elements Software package was used for post-capture image analysis. Sequential scanning was used to suppress optical crosstalk between the fluorophores in stationary-structure colocalization assays. All manipulations of contrast and illumination on color images as well as color replacement were made using Adobe Photoshop CS software (Version 23.5.5).

### 2.7 Quantitation of Phospho-Synuclein Aggregates

To quantify phospho-synuclein aggregates, 30μm floating brain sections were processed for immunofluorescent labeling with pS129 antibody. To count pS129-positive inclusions, an ImageJ macro was developed by modifying a previously described macro designed for automated cell counting.27 Briefly, the parameters of the counter were adjusted to reliably count pS129-positive inclusions. There was strong agreement between macro-counted sections and those counted manually. To count the aggregates in the macro, an area of interest was drawn on a micrograph within ImageJ, and then the macro was run to count the total number of aggregates in the outlined area. The area occupied by aggregates was manually quantified in ImageJ using the freehand tool and the measure function.

### 2.8 Unbiased Stereology

The unbiased stereological estimation of the total number of the TH+ neurons in SNpc was performed using the optical fractionator method as described previously [28]. The sampling of cells to be counted was performed using the Micro Brightfield Stereo Investigator System. The software was used to delineate the transduction area at 4x on 30 μm sections and generate counting areas of 100 x 100 μm. The estimate of the total number of neurons and coefficient of error due to the estimation was calculated according to the optical fractionator formula [28].

### 2.9 DA measurements

Frozen striatal tissue samples were shipped on dry ice to the Neurochemistry Core Lab at Vanderbilt University Medical Center, Nashville, TN for HPLC analysis of DA, 3,4-dihydroxyphenylacetic acid (DOPAC), and homovanillic acid (HVA) in the striatum. A total of 5-6 animals per group were analyzed for striatal DA metabolites.

### 2.10 Western blot Analysis

Brain tissues were collected and lysed with RIPA buffer (Cell signaling, 9806, USA) supplemented with 1% Halt Protease and phosphatase inhibitor cocktail (Thermo Fisher Scientific, 87786, USA). Samples were homogenized using a pestle (Fisherbrand™ RNase-Free Disposable Pellet Pestles; 12-141-368; Fisher, USA) and rotated at 4°C for 30 min, and centrifuged at 4°C by 12000g speed for 15 min. Based on the Bradford method, the supernatant was used for protein quantitation using a protein assay (Bio-Rad Protein Assay Kit I; 5000001; Hercules, California, USA). 80-100 μg of protein were loaded onto 4–20% Mini-PROTEAN® TGX™ Precast Gel (Bio-Rad, 4561093EDU, USA) and transferred to polyvinylidene difluoride membrane (Biorad,1704272, USA) using the Trans-Blot Turbo Transfer System (BioRad, 1704150, USA). Membranes were incubated for 1hr in 5% skim milk (Bio-Rad, 1706404, USA) prepared with 1X Tris buffered saline (Bio-Rad, 1706435, USA) with 0.1% Tween 20 (Sigma-Aldrich, P1379, USA). Primary antibodies were diluted in 5% Bovine serum albumin (Fisher, BP9703-100, USA) dissolved in 1X TBS. The following primary antibodies were used in the study: Mouse anti-αS (1:2000; BD Transduction Laboratories, Zymed Laboratories #610787), mouse anti-TH (1:2000; Millipore #MAB318), anti-hRXRα mouse (1:200; Abcam #ab50546). Secondary antibodies (1:10000 for all) were purchased from LI-COR: HRP goat anti-Rabbit IgG (926-80011, USA); HRP goat anti-Mouse IgG (926-80010, USA). Images of membranes were captured and analyzed using the Odyssey XF system (LI-COR, USA).

### 2.11 Statistical Analysis

Data were analyzed using one-way analysis of variance with Tukey post-test using Prism (GraphPad Software 10.0.2, Inc.). Data are presented as mean ± SE.

## 3. Results

### 3.1. Development of PD-like mouse model associated with αS-induced neurodegeneration

In this study, we employed an animal model of PD-like pathology, as previously described by Thakur et al and Bjorklund et al [29,30]. In order to mimic the PD pathogenesis in C57BL6 adult male mice, we used combined injections of AAV-αS and PFFs (AAV/PFF) into the SNpc. The PFFs were purchased from Stressmarq (SPR-322) and used at a concentration of 5 mg/ml. The AAV containing a blank expression cassette, which is an αS vector with an early stop codon, served as a control virus (BV) [24,31]. Monomeric αS (Stressmarq; SPR-321) was used as a control substance to PFFs. A mixture of BV and monomeric αS (BV/αS) served as a control for combined AAV/PFF administration. Mice were injected with AAVs and/or PFFs on one side of the brain; the other side was kept as a non-transduced internal control (Figure 1).

**Figure 1.**
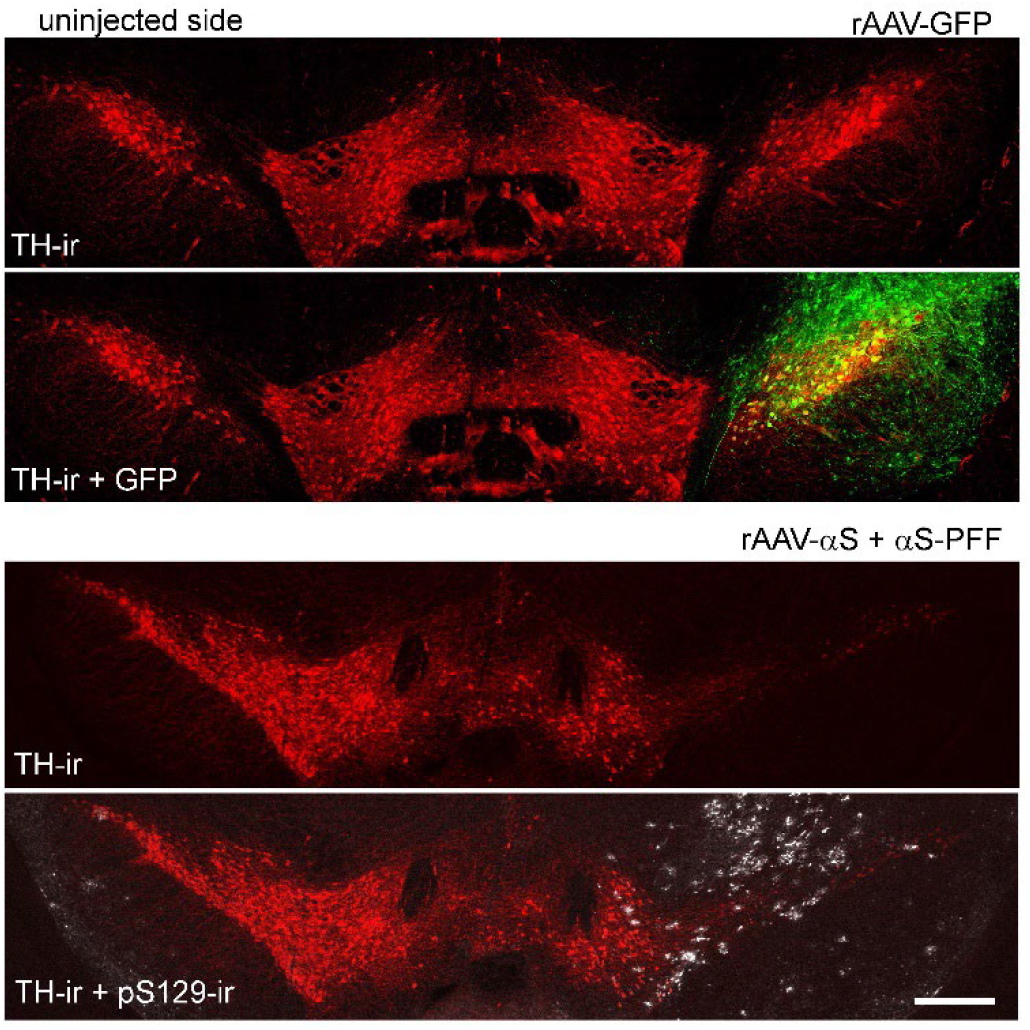
Fluorescent microscopy of the mouse SNpc at 8 weeks after AAV-GFP and AAV/PFF injections. The upper image shows the expression of the GFP transgene on the injected side of mouse brain. GFP (green) was expressed in most TH+ neurons (red) in SNpc and also can be seen in the SN pars reticulata, as well as surrounding neuronal structures of the mesencephalon. The bottom image illustrates pS129-positive inclusions (white) in SNpc neurons and nearby structures. Scale bar: 0.5 mm.

The study involved several groups of animals: uninjected controls, and those injected with single AAV-αS, BV, AAV-GFP, AAV-RXR, and a combination with AAV/PFF or BV/αS. We established injection conditions to secure αS expression level and consistent with the reported αS levels in postmortem human PD brains32, we used the AAV-αS titer that induced moderate levels of αS overexpression, which was three-fold higher relative to endogenous levels. To confirm this, we first validated a mouse anti-αS antibody capable of recognizing an epitope shared by both endogenous mouse αS and exogenous human αS (Figure 2A). In these experiments, the ratio of αS in the injected versus uninjected mouse SNpc was measured at 4 weeks post-injection, a time point when neurodegeneration was not yet significant. The αS ratio varied between 2.7 and 3.1, with an average increase of 2.9-fold in total αS expression (Figure 2B).

**Figure 2.**
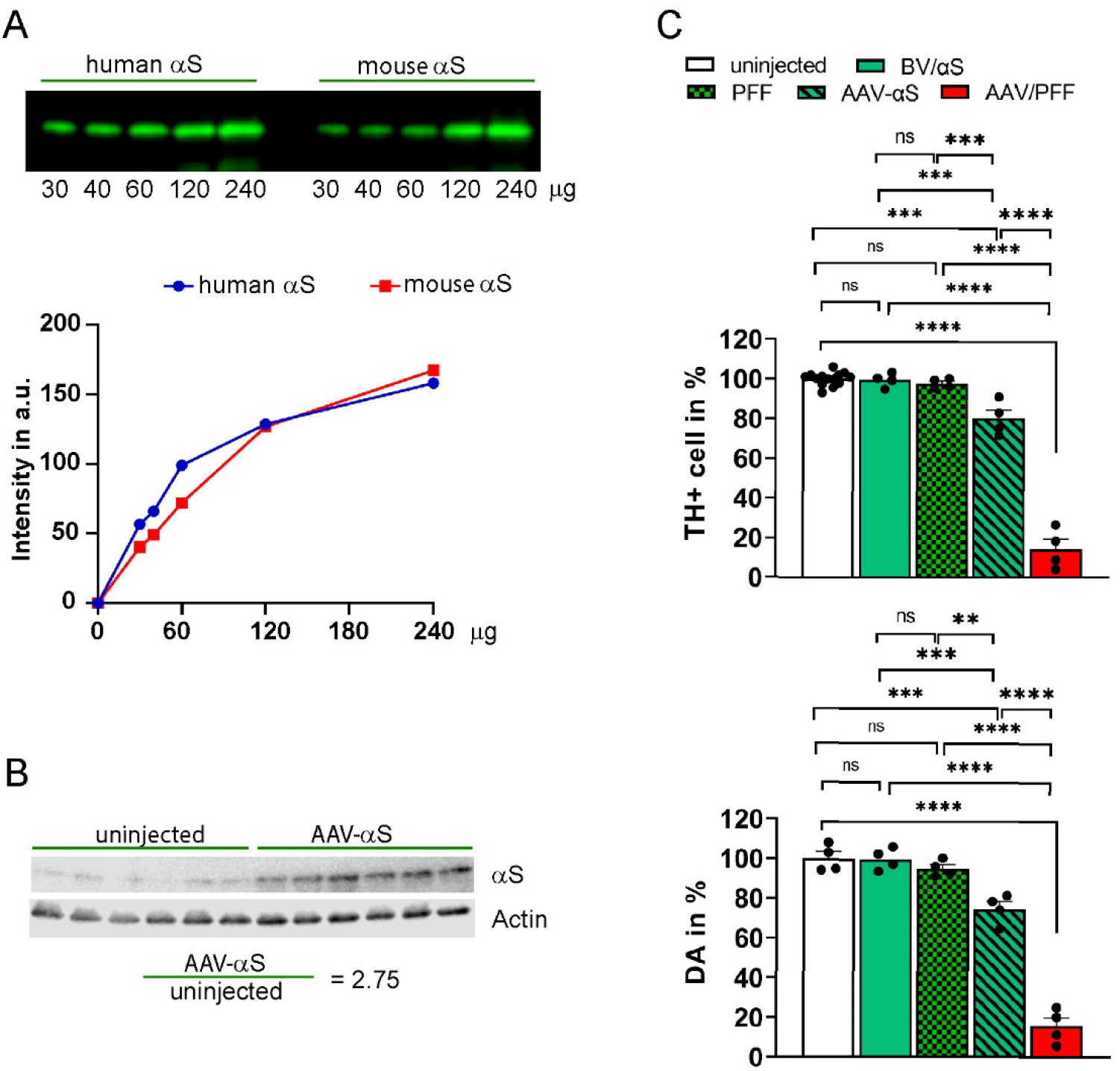
Comparative analysis of αS mouse models of PD. All AAV vectors have the same volume and titer (6 x1012 vg/ml). The total injection volume (2 μl) of AAV-GFP and AAV-αS was adjusted using PBS to equalize with BV/αS or AAV/PFF. (A) The comparative affinity test of anti-αS antibody used in the study to bind recombinant human and mouse αS proteins of known concentration at 4 weeks post-injection. (B) Measurement of αS expression in animals that had been injected with human αS at 4 weeks postinjection. Western blot (WB) showing nigral extracts taken from the uninjected and injected sides. Thirty micrograms of tissue extract was analyzed by WB using an antibody that recognize both human and mouse αS. The amount of αS was normalized to β-actin and the ratio of the injected versus uninjected sides was calculated (n=6). (C) Combined AAV/PFF administration into SNpc resulted in more severe loss of nigral TH+ cells and striatal DA compared with individual injections of AAV-αS or PFF at 8 weeks post injection. Upper graph demonstrates unbiased estimation of nigral TH+ cells in SNpc of mice injected with AAV-αS or PFF and a combination of AAVs/PFF or BV/αS. Tissue slices were labeled with antibody to TH as described in Materials and Methods and the percentage of surviving cells was calculated by comparison with the uninjected side in the same animal. Lower graph shows measurement of striatal DA. The amount of DA in striatal tissue was measured on the injected and uninjected sides of individual animals and displayed as the mean percentage of DA remaining on the injected side compared with the uninjected side plus standard error. Both graphs demonstrate the significant loss of nigral TH+ cells and striatal DA in AAV/PFF injected mice. p*< 0.05; **<0.01; ***<0.001; ***,0.0001 (n=4 for each group except of TH+ cell count in uninjected control, n=14).

We observed significant striatal dopamine (DA) and TH+ neuronal loss at 8 weeks post-injection. These effects were most dramatic in mice injected with AAV/PFF as compared to those receiving either rAAV-αS or PFFs alone (Figure 2C), highlighting the advantages of the combined approach in accelerating the development of LB-like features. Notably, no differences were observed between uninjected mice and BV/αS injected controls, nor between control groups and PFFs injections alone. However, substantial differences were detected between control groups and AAV-αS as well as between control groups or AAV-αS and AAV/PFF (Figure 2C). As a result of AAV/PFF injections, we observed the formation of pS129-positive inclusions (Figure 1). These inclusions were associated with increased neuronal vulnerability seen as dramatic reduction in number of TH+ cells in the SNpc accompanied by a decline in striatal DA levels (Figure 2C).

### 3.2. Combined AAV/PFF injection induce severe neurodegeneration accompanied by inflammatory response and gliosis

To investigate a link between the observed aggregates, TH+ cell loss, and inflammation, we next analyze the level of GFAP. GFAP is upregulated in response to CNS injury or disease and serves as a marker of astrogliosis. Elevated GFAP levels are observed in conditions such as AD, PD, HD, and ALS and contribute to neurodegeneration via pro-inflammatory signaling [33-35]. During activation, accompanied by upregulated GFAP expression, astrocytes release pro-inflammatory cytokines and chemokines (e.g., IL-1β, TNF-α), which promote immune cell recruitment. Another protein, IBA1, serves as a marker for microglia and their activation, reflecting microglial dynamics in CNS health and disease. Its upregulation in activated microglia makes it a critical tool to study neuroinflammation. Activated microglia release cytokines such as IL-1β, TNF-α, and IL-6, which can exacerbate neuroinflammation. Therefore, we analyzed these markers in the PD brains.

Figure 3 demonstrates a significant increase in both GFAP and IBA1 fluorescent signals ranging from 2-to 3-fold in AAV/PFF injected brains previously identified with dramatic TH+ cell loss compared to BV/αS injections. Results of this experiment indicated a notable inflammatory response associated with the combined AAV/PFF approach.

**Figure 3.**
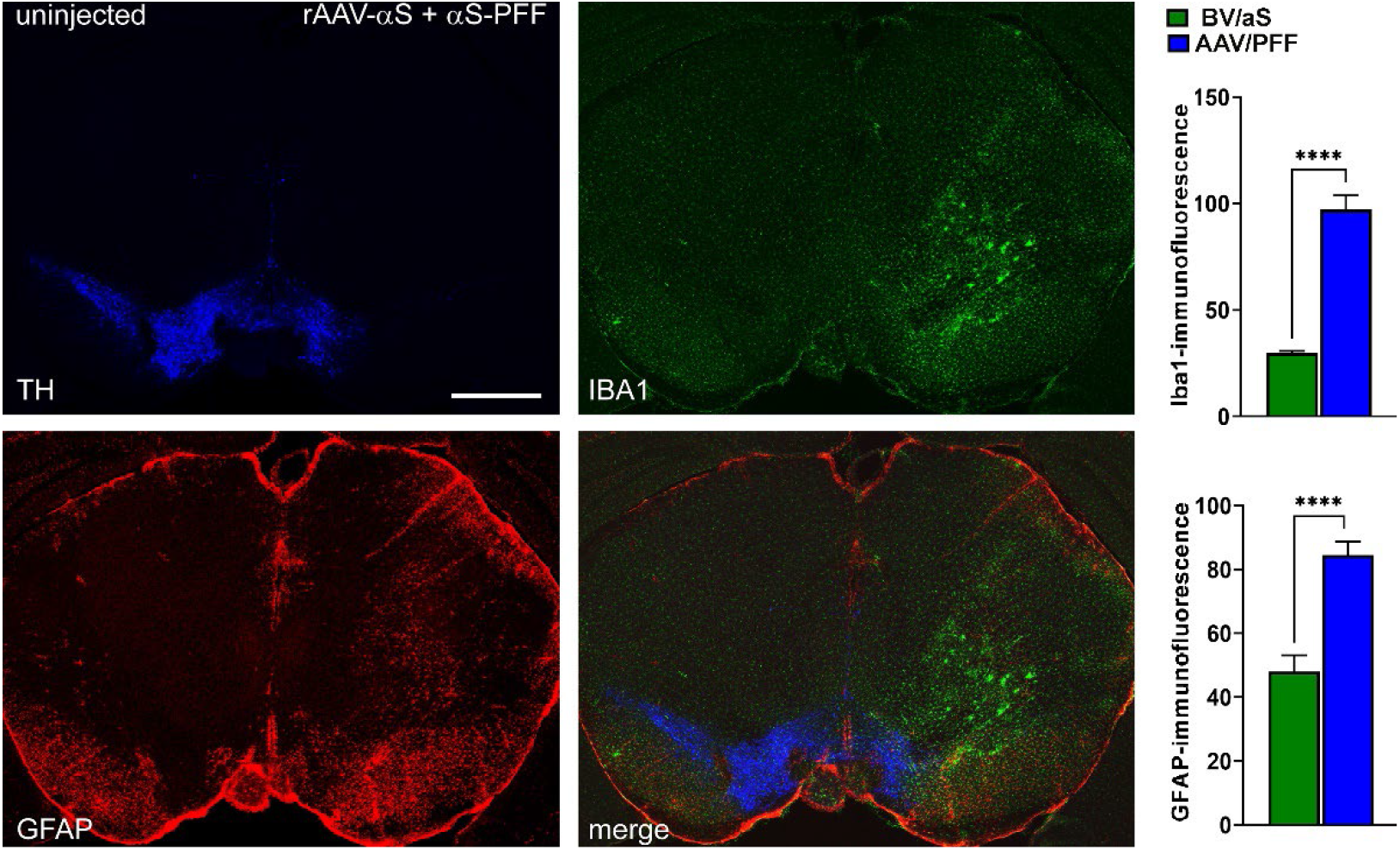
Combined AAV/PFF injection led to significant inflammation response in mouse brain evident by increase in GFAP and IBA1, markers of astrocytes and microglia. Of note, we did not find the difference between uninjected and injected SNpc of any control groups. p< ***,0.0001 (n=5-6). Scale bar: 1mm.

### 3.3. RXR diminishes LB-like formation in primary mouse cortical neurons

Preliminary testing of AAV-RXR vector in primary mouse neuronal cells (Figure 4) demonstrates a powerful RXR effect on the formation of 129-positive LB-like inclusion. Despite diffuse pS129 immunostaining was observed in both GFP and RXR experimental groups, LB-like αS inclusions (denoted by arrows) localized to cell bodies were identified only in GFP-positive cells after nine days of exposure to PFFs and monomeric αS. None of these inclusions were found in hRXRα-expressing neurons, suggesting the therapeutic potential of treatment with exogenous RXR.

**Figure 4.**
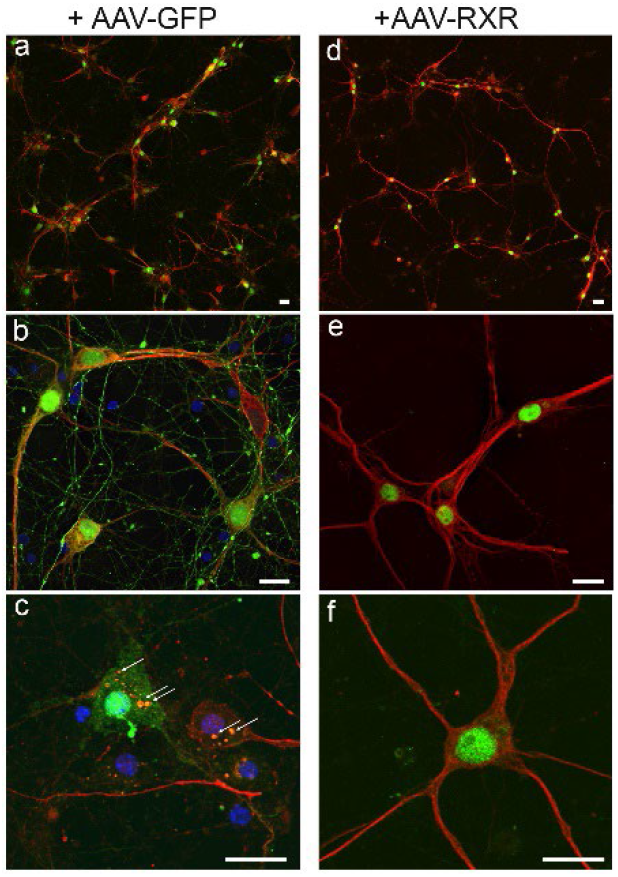
Confocal images illustrate AAV-mediated expression of native GFP (a-c) and human RXRα (d-f), both shown in green, in cortical neurons at DIV14 after nine days of exposure to PFFs and monomeric αS. Both experimental groups demonstrate immunofluorescent staining with anti-pS129 antibody (shown in red). However, LB-like αS inclusions localized to cell bodies denoted by arrows were identified only in GFP-positive cells and none of them were found in hRXRα expressing neurons. DAPI staining (in blue) added in (c). Scale bars: 25 μm.

### 3.4. RXR overexpression significantly ameliorate PD-associated pathology in mouse PD model

Previous studies indicated that reduced RXR signaling triggers neuronal stress and neuroinflammation in PD pathobiology [6,8]. To explore the therapeutic potential of RXR activation for PD treatment, we utilized AAV-mediated transduction to deliver either human RXRα or control GFP, alongside AAV/PFF, into the mouse SNpc. Various control and experimental groups were included in the study, incorporating combinations of GFP, human RXRα viral vectors, and control BV/αS. The mice were analyzed at 4-and 8-weeks post-injection (Figure 5 and Figure 6). An unbiased stereology count of nigral TH+ cells in the injected SNpc was compared with the uninjected SNpc for each mouse. No significant differences were observed between animals injected with the GFP, human RXRα vectors, and/or control BV/αS, and the uninjected side at both the 4- and 8-week time points (p = 0.963; data not shown). Therefore, the BV/αS control group at both time points was used for further comparisons.

**Figure 5.**
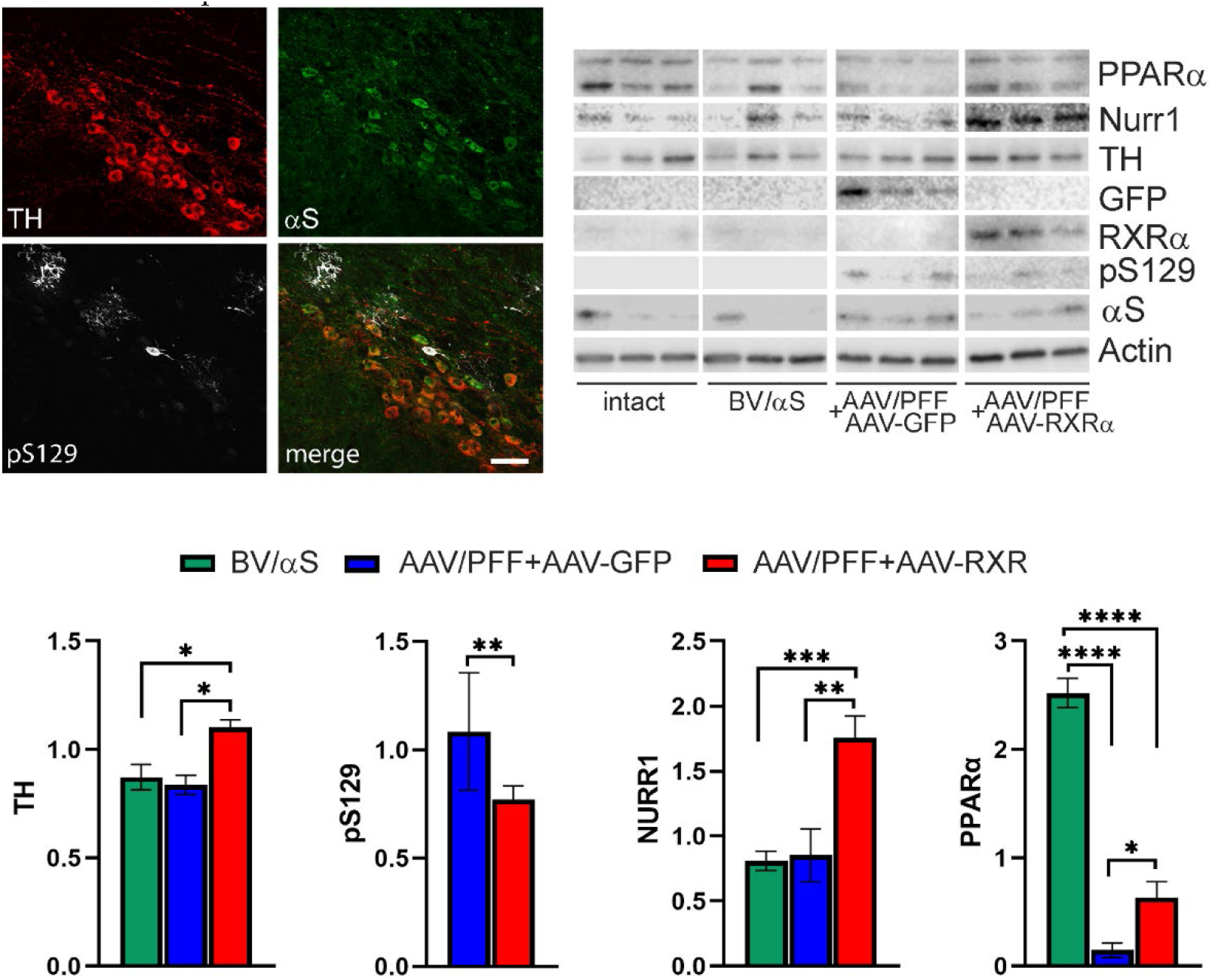
The SNpc of mice at 4-week after PFFs and AAV vector injections. Confocal imaging reveals pS129-positive aggregates in TH+ neurons in SNpc of mice treated with AAV/PFF. WB analyses show no difference in nigral TH protein levels between control groups and AAV/PFF + AAV-GFP injected brains, which correlates with stereology counts of TH+ cells in this time point. However, TH protein level was increased in AAV/PFF + AAV-RXR treated nigras. The levels of PPAR and NURR1 were significantly downregulated in AAV/PFF injected brains compared to BV/αS and AAV/PFF + AAV-RXR injected brains. No difference was observed between BV/αS and uninjected brains. The level of pS129 protein adjusted to total αS was significantly downregulated in AAV/PFF+AAV-RXR compared to AAV-PFF+AAV-GFP mice. p*< 0.05; **<0.01 (n=6). Scale bar: 25 μm.

**Figure 6.**
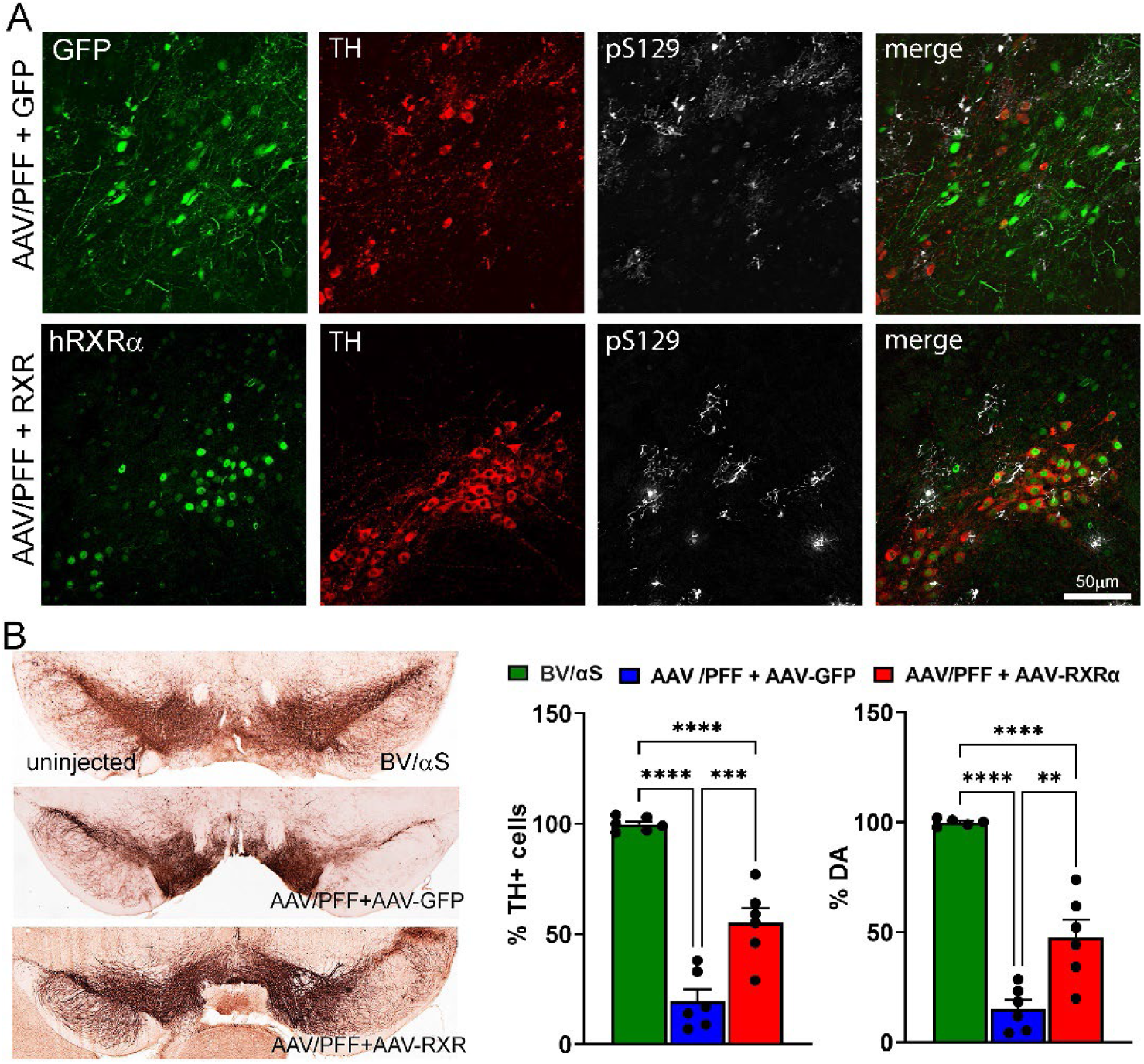
AAV-mediated overexpression of human (h) RXRα ameliorates PD-like pathology induced by AAV/PFF intra-nigral injection. Confocal imaging demonstrates expressions of transgenic GFP or hRXR (in green) within TH+ neurons (in red) with pS129 deposits (in white). Unbiased stereology counts of TH+ cells and measurement levels of striatal DA are shown. p*< 0.05; **<0.01; ***<0.001; ****<0.0001.(n=6).

Figure 5 presents the study results of the mouse SNpc terminated at 4 weeks post-injection with AAV/PFF. Despite confocal imaging revealing pS129-positive inclusions in TH+ neurons in the SNpc and surrounding nervous structures of mice treated with AAV/PFF, no significant nigral TH+ cell loss or marked reduction in striatal DA were detected in the mouse brains at this point. However, a notable reduction in PPARα levels was observed in the nigral tissue of AAV/PFF-injected mice at 4 weeks post-injection. At the same time, co-injection of AAV/PFF with AAV-RXR led to over 2-fold increase in PPARα level compared with AAV/PFF + AAV-GFP injected mice. In addition to PPARα, RXR overexpression in AAV/PFF-injected mice enhanced NURR1 production in the SNpc at 4 weeks post-injection. This implies that RXRα overexpression in the PD-like brain activates NURR1, which has been proposed as a standalone therapeutic target for PD [22] Consistent with the previous study demonstrating NURR1-induced TH production [17], this NURR1 upregulation was accompanied with significant increase in TH protein. Moreover, the level of pS129 protein was significantly downregulated (*p*<0.001) in AAV/PFF+AAV-RXR compared to AAV-PFF+AAV-GFP mice at 4 weeks postinjection. This finding indicates that NURR1 is a downstream target of RXR activation and underscores the potential of RXR-mediated NURR1 therapy.

At 8 weeks post-injection, while AAV/PFF treatment induced a significant loss of nigral TH+ cells and striatal DA, the simultaneous delivery of human RXRα and AAV-PFF diminished TH+ cell loss and increased DA level (Figure 6). Additionally, RXRα overexpression led to a decrease in GFAP immunoreactivity and GFAP+ punctata, indicating reduced recruitment of astrocytes to the AAV/PFF-affected area at 8 weeks post-injection (Figure 7). It is also noteworthy to mention that the microglial marker IBA1 was diminished in RXR-overexpressing brains with pS129-positive inclusions, further supporting the anti-inflammatory effects of RXRα overexpression.

**Figure 7.**
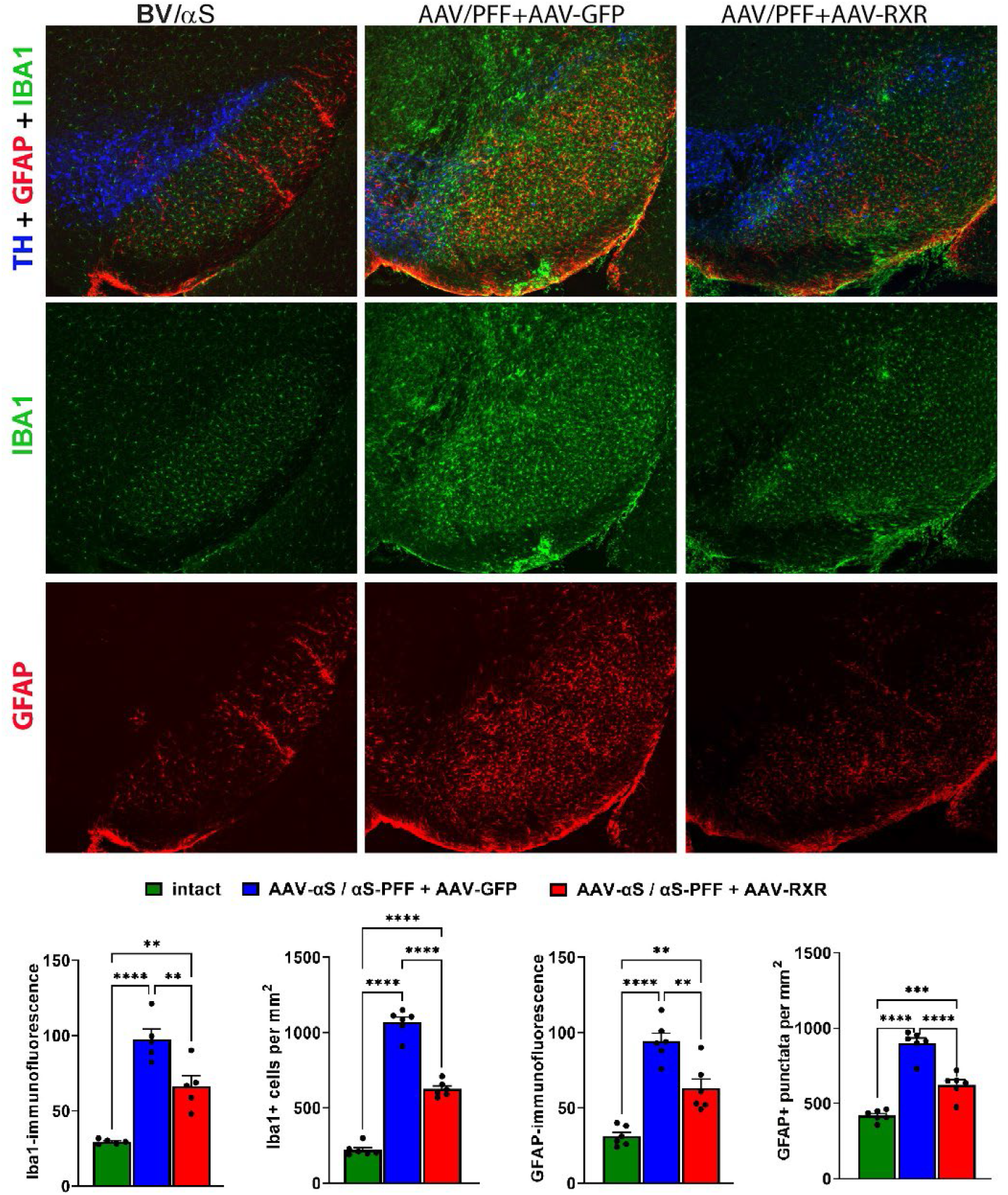
Confocal images illustrate reduced expression of inflammatory markers, GFAP (in red) and IBA1 (in green), in AAV/PFF brains upon AAV-RXR delivery. TH+ cells are shown in blue. The graphs depict the results of immunofluorescent detection of GFAP and IBA1 expression shown in arbitrary units. p*< 0.05; **<0.01; ***<0.001; ****<0.0001.(n=6).

Motor behavior impairment was assessed using the cylinder test to measure asymmetry in forelimb use. At 4 weeks post-injection, no significant differences were observed between experimental groups (Figure S1). However, by 8 weeks post-injection, the AAV/PF + AAV-GFP group exhibited significant motor impairment compared to the control BV/αS group. In contrast, mice injected with AAV/PFF + AAV-RXR showed a trend toward improvement relative to the AAV/PF + AAV-GFP group, though this difference did not reach statistical significance (Figure S1).

Finally, we analyzed the distribution area of pS129-positive inclusions affected by RXR overexpression by comparing injection sites of AAV/PFF + AAV-GFP and AAV/PFF + AAV-RXR mice. We counted the pS129-positive inclusions in outlined areas (Figure 8). Remarkably, the number of aggregates and the size of the area containing pS129+-LB-like aggregates, measured in μm^2^, significantly decreased with the distance from the point of injection in AAV/PFF + AAV-RXR injected brains compared to AAV/PFF + AAV-GFP injections (p<0.0001).

**Figure 8.**
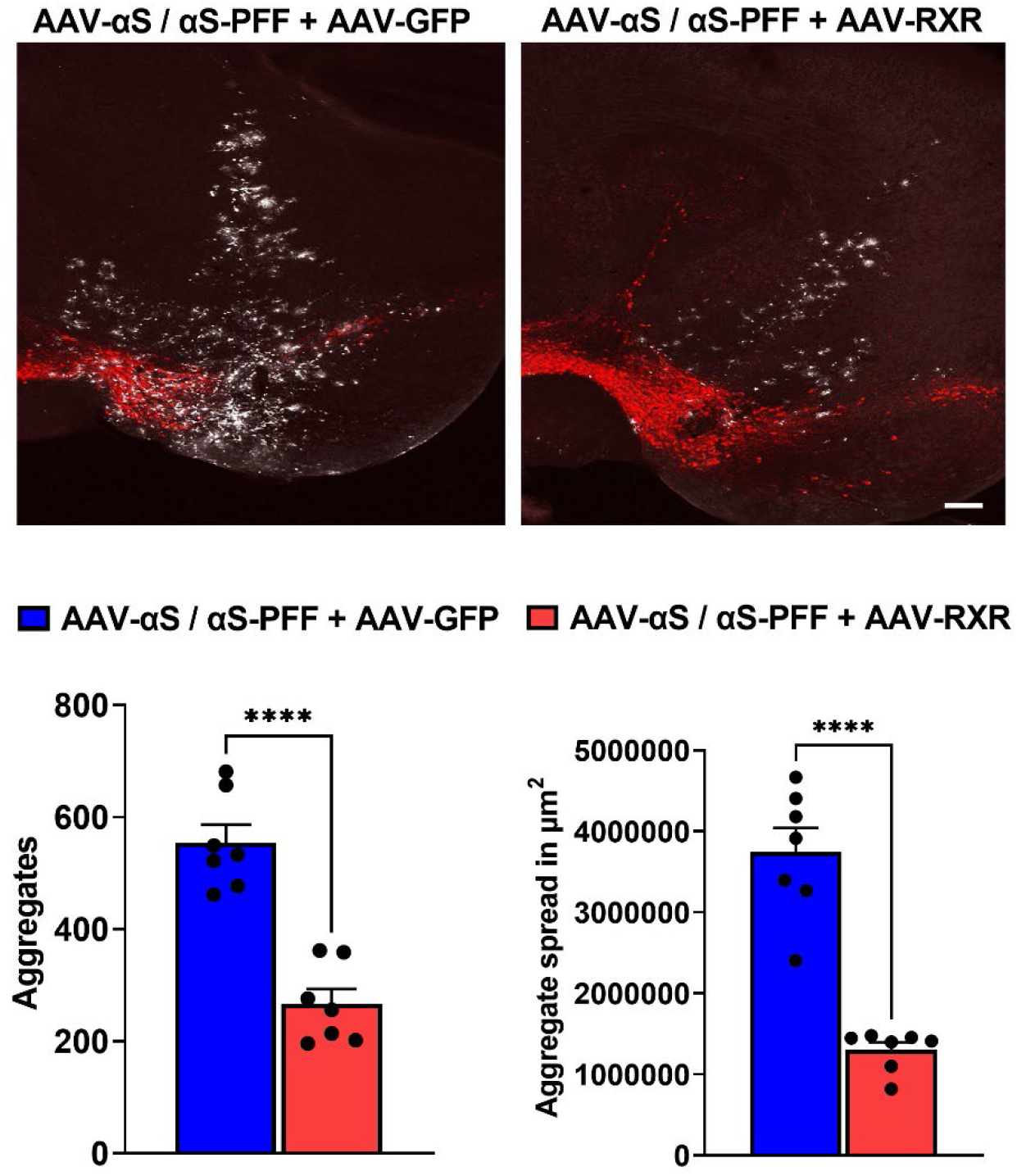
The pS129-positive inclusions (white) at injection sites of mouse brains treated with AAV/PFF + AAV-GFP and AAV/PFF + AAV-RXR. TH-positive cells stained in red. Graphs represent comparative count and distribution area of pS129-positive aggregates. Student t-test, p****<0.0001.

## 4. Discussion

In the current study, we present a reliable mouse model that mimics PD-like pathology, including the accumulation of LB-like aggregates, TH neuronal loss, DA deficit, and activation of astrocytes and microglia. Our data indicate that a combination approach delivering AAV-αS and commercial PFFs to the SNpc results in the activation of a severe inflammatory response. This approach significantly accelerates, and mimics PD progression compared to previously reported models based on individual AAV-αS or PFFs injections. This confirms the observation by Bjorklund A et al [30] that the combined AAV/PFF delivery offers several advantages over the standard PFF model due to the enhanced and accelerated αS pathology and glial response induced by the PFFs and elevated αS levels, as shown in Figures 2C and 3. An advantage of the current study is that we demonstrated that the AAV/PFF-mediated neurodegeneration provides a window of opportunity for drug delivery according to our data. A limitation of the current study is that the RXR-based intervention in the PD pathogenesis should be tested at later stages with marked disease progression thus mimicking more of the real-life situation in individuals with PD.

The AAV/PFF mouse model provides benefits by triggering an early inflammatory response observed at 3-4 weeks post-injection before severe aggregate formation is observed, as previously reported [30]. Notably, the significant striatal DA and nigral TH+ cell loss was not yet detected at this time point; however, a level of protein expression induced by AAV-mediated gene transfer reaches a maximum [25,36,37]. This point allows the study pathogenic mechanisms triggered by AAV/PFF.

Our data indicates that overexpression of human RXRα in the SNpc ameliorates αS-associated neurodegeneration. This was evident through a diminished loss of striatal DA and nigral TH+ cells, and reduced deposition of pS129-positive aggregates. Furthermore, we revealed reduced GFAP and IBA1-positive fluorescent signaling and number of positive cells. Given the limitation of the current study, including lack of the detailed analysis of activated astrocytes and microglia phenotypes, this reduction suggests fewer recruited astrocytes and IBA1-positive microglial cells, overall indicating a mitigation of neuroinflammation. In agreement with diminished neuroinflammation, the reduction in αS deposits in the PD brain also suggests that RXR upregulation prohibits the aggregation of an insoluble form of the protein. The results of TH+ cell counting, and striatal DA assay align, indicating the neuroprotective effects of RXR upregulation.

Based on our data, we anticipate that PPAR is necessary for RXR-based therapies to enhance TH cell survival and DA levels in the PD brain. We revealed that AAV/PFF treatment downregulates PPARα production. In turn, RXR overexpression induces significant upregulation of PPARα. It is well known that PPAR α, β, and γ are the permissive RXR binding partners; when activated in the central nervous system, RXR plays a critical role [9]. Recent studies have revealed reduced PPARα expression in AD brains [38] and the genetic linkage to the disease [39,40]. The neuroprotective effect of PPAR agonists have recently been proposed using neurological murine models and include anti-inflammatory effects, APP degradation, and Aβ inhibitory functions [41-45] The pharmacological modulation of PPARs by pioglitazone, rosiglitazone, ibuprofen, piroxicam, ciglitazone, and GW1929 was shown to be neuroprotective in AD, ALS, MS, and PD models [46-51]. In particular, in MPTP and 6-OHDA models of PD, preclinical studies have shown that different PPARγ agonists reduce neuronal cell death, microglial activation, and DA level in the hippocampus and the SNpc. [52] [19,53] [15,54]. A conformational change in the RXRa/PPARγ complex allows the heterodimer to bind PPAR-response elements (PPRE) in the nucleus to activate gene transcription. Therefore, it is not surprising that our results have demonstrated that the AAV-mediated overexpression of human RXRα increases the expression of PPARα in mouse brains with PD-like pathology.

Our data also indicate that in addition to PPARα, RXR overexpression in mice with experimental α-synucleinopathy enhances NURR1 production in SNpc. This finding is in accordance with the multiple studies that highlighted the significance of changes in NURR1 expression during the pathogenesis of PD and related disorders. Thus, it has been shown that NURR1 is significantly decreased in the PD nigral neurons containing αS-immunoreactive inclusions and in the DA nigral neurons with neurofibrillary tangles, correlating with the loss of TH and diminished intracellular pathology in both αSNP and tauopathies [55]. Furthermore, downregulation of NURR1 promotes upregulation of the NF-κB/ NLRP3 inflammasome axis [56-58], while NURR1 overexpression and activation by means of HX600 agonist treatment have significant anti-inflammatory effects decreasing the expression of genes involved in inflammation control [59] and reducing microglia-expressed proinflammatory mediators, preventing neuroinflammation and neuronal cell death [17]. It is also worth mentioning that the TH upregulation and anti-inflammatory therapeutic effect from the application of HX600 agonist is associated with the activation of the NURR1/RXR complex, suggesting their dual role in PD development.

In summary, the current study not only validated a PD-like mouse model using commercially available PFFs, making it convenient to study the role of genetic modifiers, but also provided evidence that mice with progressive αSNPs present a model for druggable intervention in PD progression. Our study demonstrated the power of RXRα-based therapy to mitigate PD progression by targeting major pathological events, such as the formation of LB-like structures, loss of TH+ neurons, and neuroinflammatory response. The mechanism of RXR-based intervention requires future investigation. At this point, it is challenging to anticipate whether enhanced RXR will demonstrate a preference for binding to one nuclear receptor over another. Moreover, the RXR-based therapy may enhance the pool of RXR homodimers available to partner with both NURR1 and PPARs to regulate the expression of genes responsible for anti-inflammatory action.

## Institutional Review Board Statement

The animal study protocol was approved by the University of Alabama at Birmingham - Institutional animal care and use committee (protocol no. 10085).

## Data Availability Statement

The research data is available upon request. The raw data are also available as a supplemental material.

## Acknowledgments

We would like to thank Dr. David S. Standaert for his critical evaluation of the manuscript and for providing valuable insights that enhanced its quality.

---

**Figure S1.**
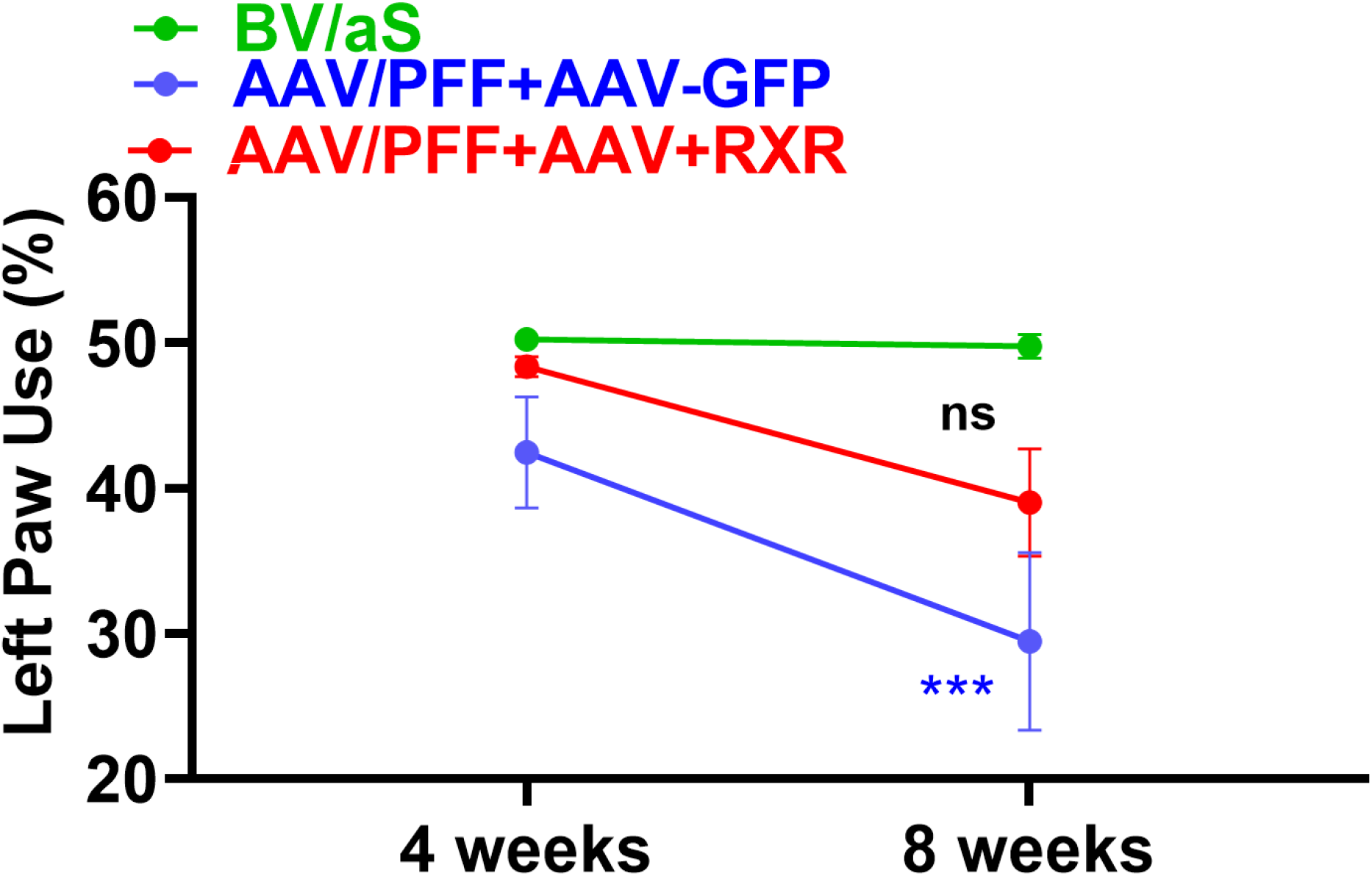
Cylinder test.

